# HIV-1 Vpr Degrades TET2 to Suppress IRF7 and Interferon Expression in Plasmacytoid Dendritic Cells

**DOI:** 10.1101/823328

**Authors:** Qi Wang, Li-Chung Tsao, Lei Lv, Yanping Xu, Liang Cheng, Haisheng Yu, Liguo Zhang, Yue Xiong, Lishan Su

**Affiliations:** Lineberger Comprehensive Cancer Center, University of North Carolina at Chapel Hill, North Carolina, USA; Department of Microbiology & Immunology, University of North Carolina at Chapel Hill, North Carolina, USA; IBP-CAS, Beijing, China

**Author notes:** Correspondence: Lishan Su.

**Keywords:** HIV-1, Vpr, TET2, pDC, IRF7, IFN-I

## Abstract

Plasmacytoid dendritic cells (pDCs) are the major source of type I interferons (IFN-I) in rapid response to viral infections, with constitutive expression of interferon regulatory factor 7 (IRF7). HIV-1 expresses several accessory proteins to counteract specific IFN-induced host restriction factors. As one abundant virion-associated protein, HIV-1 Vpr remains enigmatic in enhancing HIV-1 infection via unclear mechanisms. Here we report that Vpr impaired IFN-I induction in pDCs to enhance HIV-1 replication in CD4+ T cells. Blockade of IFN-I signaling abrogated the effect of Vpr on HIV-1 replication. Virion-associated Vpr suppressed IFN-I induction in pDC by TLR7 agonists. Modulation of IFN-I induction by Vpr was genetically dependent on its activity of TET2 degradation. We further demonstrate that Vpr-mediated TET2 degradation reduced expression of IRF7 in pDCs. Finally, degradation of TET2 in pDCs by Vpr reduced the demethylation level of the IRF7 promoter via CXXC5-dependent recruitment. We conclude that HIV-1 Vpr functions to promote HIV-1 replication by suppressing TET2-dependent IRF7 expression and IFN-I induction in pDCs. The Vpr-TET2-IRF7 axis provides a novel therapeutic target to control HIV-1 infection.

## Introduction

During human immunodeficiency virus type 1 (HIV-1) infection, plasmacytoid dendritic cells (pDCs) are the major type I interferons (IFN-I) producer cells (Liu, 2005). The CD4+ pDCs are rapidly activated and are involved in initiating antiviral immunity and in suppressing HIV-1 replication.

To sense pathogen-associated molecule patterns (PAMPs) such as viral RNA or DNA, pDCs express the endosomal Toll-like receptors TLR7 and TLR9, which bind single-stranded RNA and CpG-containing DNA, respectively (Diebold et al., 2004; Hemmi et al., 2000). Activation of TLR7 or TLR9 leads to activation of MyD88 and subsequent phosphorylation and nuclear translocation of IFN regulatory factor 7 (IRF7) to bind the IFN-I promoter in pDCs (Guiducci et al., 2008; Honda et al., 2005b), which is critical for IFN-I induction in pDCs (Honda et al., 2005a; Smith et al., 2016). In contrast to other cell types, IRF7 is constitutively expressed in pDC and its activity in pDCs is regulated by both transcriptional and posttranscriptional mechanisms (Colina et al., 2008; Kim et al., 2016; Lee et al., 2013; Ma et al., 2017). Interestingly, the promoter of IRF7 contains methylated CpG islands that can be demethylated by the DNA demethylase TET2 via CXXC5-dependent recruitment (Ma et al., 2017).

Although HIV-1 can infect pDCs in humanized mice and human patients (Donaghy et al., 2003; Zhang et al., 2011), pDCs are not efficiently infected by HIV-1 in vitro. However, HIV-1 or HIV-infected cells can activate pDCs to produce IFN-I via CD4-dependent endocytosis without fusion-dependent productive infection (Beignon et al., 2005). In spite of rapid induction of IFN-I, HIV-1 persists and establishes chronic infection in the infected hosts. To counteract IFN-induced anti-viral immunity, HIV-1 expresses several accessory proteins to target specific IFN-stimulated genes (ISGs) or host restriction factors to maintain persistent replication. For example, HIV-1 Vif targets the ISGs APOBEC3 proteins to protect its genome integrity during replication and Vpu targets the ISG tetherin for degradation to allow release of HIV-1 virions from producer cells. The SIV Vpx is reported to target the SAMHD1 protein to facilitate SIV replication in macrophages and dendritic cells.

As one abundant virion-associated protein, however, HIV-1 Vpr remains enigmatic in enhancing HIV-1 infection via unclear mechanisms. It has been reported that, in nonhuman primates infected with Vpr mutant SIV, the Vpr mutations are reverted, resulting in a functional *Vpr* gene over the course of infection (Goh et al., 1998; Lang et al., 1993). Monkeys infected with a Vpr-null SIV mutant showed decreased virus replication and delayed disease progression (Gibbs et al., 1995; Hoch et al., 1995) and mutant Vpr in HIV-1 delayed viral replication in humanized mice (Sato et al., 2013). HIV-1 Vpr is a 14 kDa protein that is efficiently packaged in HIV-1 virions (Selig et al., 1999), reportedly with 250-700 copies of Vpr proteins per HIV-1 particle (Muller et al., 2000). Upon HIV-1 entry, Vpr is released in target cells, which is implicated in enhancing early steps of viral replication and/or immune evasion. Biochemically, Vpr has been reported to interact with VprBP to mediate several key cellular activities including G2 cell cycle arrest (Wen et al., 2007). In a recent report, Vpr was postulated to cause G2 arrest and help escape immune sensing by activation of the SLX4 complex in target cells (Laguette et al., 2014), although SLX4-independent mechanisms are reported (Fregoso and Emerman, 2016; Zhou et al., 2016). In addition, Vpr has also been reported to target IRF3 for degradation in PM1 cells for immune evasion(Okumura et al., 2008) and to block the phosphorylation of TBK1 in myeloid cells to reduce IFN production (Harman et al., 2015). We have recently reported that Vpr/VprBP-CLR4 binds and degrades the DNA demethylase TET2 to enhance HIV-1 replication in macrophages via its effect on IL6 resolution(Lv et al., 2018) and IFITM3 expression(Wang and Su, 2019). The identification of the human hub silencing (HUSH) complex as a target for Vpr-mediated degradation also suggests that HIV-1 Vpr may alter chromatin remodeling complexes and host responses(Yurkovetskiy et al., 2018).

We extended our study and found that Vpr suppressed type 1 interferon induction in pDCs to enhance HIV-1 replication in co-cultured CD4+ T cells. In addition, we discovered that Vpr impaired IFN-I induction in pDCs through its binding with VprBP/TET2 degradation but independent of its G2 arrest activity. Lastly, Vpr led to reduced IRF7 expression by preventing its TET2-dependent demethylation. These findings reveal Vpr as the first HIV-1 protein that suppress host interferon induction and suggest targeting the Vpr-TET2-IRF7 axis in pDCs may prove clinically beneficial to control HIV-1 infection.

## Results

### Vpr enhances HIV-1 replication in CD4 T cells co-cultured with pDCs by attenuating IFN-I induction in pDCs

CD4+ T cells are the major target cells for HIV-1 replication (Perelson et al., 1993), and Vpr doesn’t affect HIV-1 replication in activated CD4+ T cells (Balliet et al., 1994; Planelles et al., 1995). Cell-free HIV-1 particles are poorly detected by pDCs, but HIV-1-infected CD4^+^ T cells are effectively sensed by pDCs to induce IFN-I expression (Colonna et al., 2004),(Lepelley et al., 2011). HIV-1 or HIV-1ΔVpr infection in purified CD4+ T cells led to similar levels of HIV-1 infection (Figure 1A and S1). To determine if Vpr affects pDCs to modulate HIV-1 replication in CD4+ T cells, we co-cultured purified pDCs (Figure 1B and S1) with autologous CD4 T cells already infected with wild type HIV-1 or HIV-1ΔVpr viruses. As shown in the Figure 1C, p24 levels from HIV-1 or HIV-1ΔVpr viruses-infected cells are comparable in the absence of pDCs. We also showed that, in the co-culture system, p24 was predominantly from infected CD4+ T cells but not pDCs (Figure S1C). However, CD4+ T cells infected with HIV-1ΔVpr mutant viruses were more sensitive to co-cultured pDCs (Fig. 1C/D). Summarized data from three different donors showed that Vpr could resist ∼50% of pDCs-mediated inhibition of viral replication. correlated with lower IFN-α induction (Fig. 1E). CD4+ T cells infected with HIV-1 or HIV-1ΔVpr viruses, or pDC alone, showed undetectable IFN-α induction (Fig. 1E). To determine whether the Vpr effect on HIV-1 replication was due to different IFN-I induction, we utilized an antibody against human IFN- α/β receptor 1 (α-IFNAR1) to specifically block IFN-I signaling (Cheng et al., 2017). Consistently, the IFNAR blocking mAb enhanced HIV-1 replication in CD4 T cells infected with HIV-1 or HIV-1ΔVpr (Fig. 1F), and abrogated the Vpr activity to further enhance HIV-1 replication in CD4+ T cells cocultured with pDCs (Fig. 1F)(Kerkmann et al., 2003).

**Figure 1.**
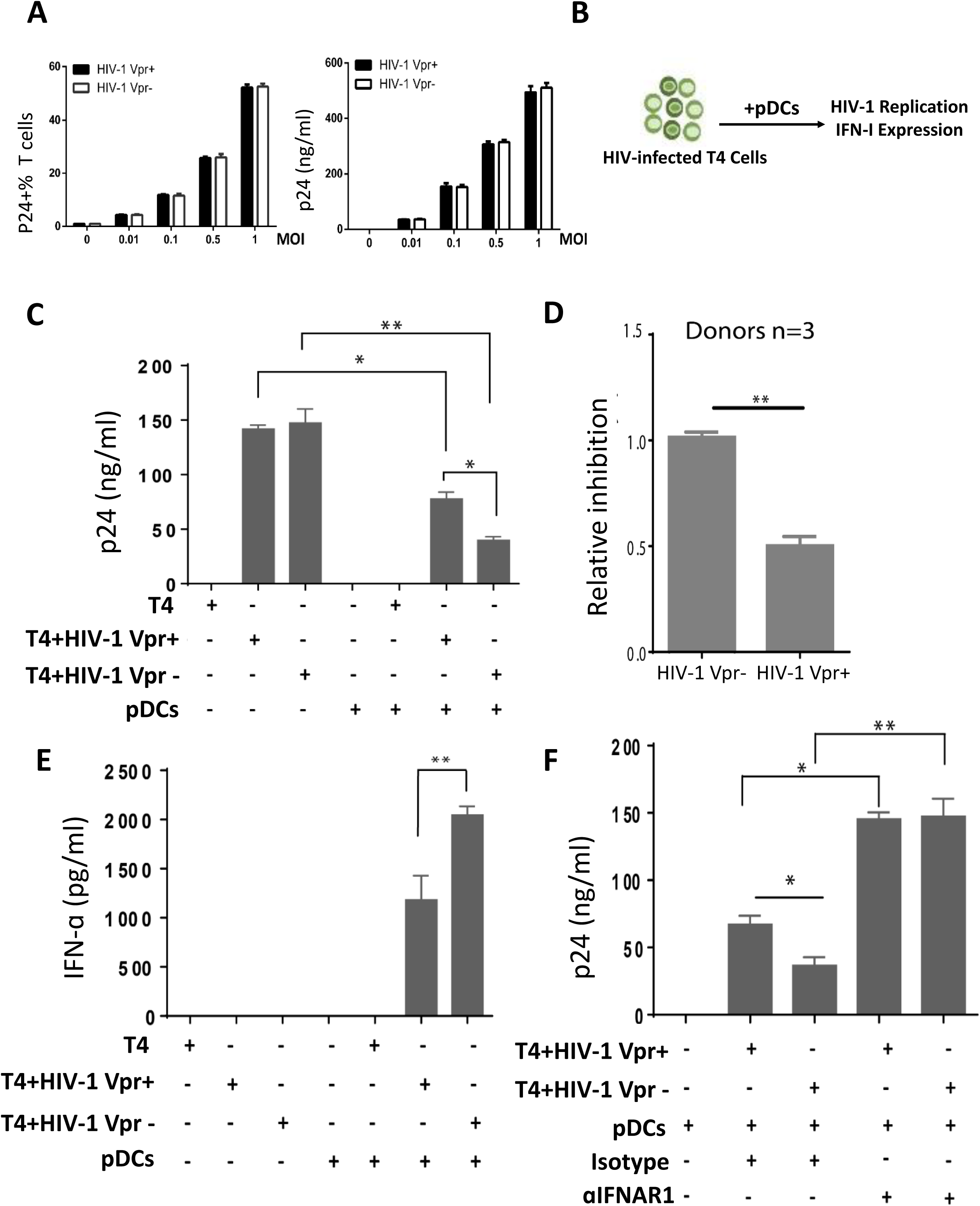
Vpr inhibits IFN-I induction in pDCs to enhance HIV-1 replication in co-cultured CD4+ T cells. (A) HIV-1 or HIV-1ΔVpr viruses infect and replicate in activated CD4+ T cells at similar levels. HIV p24+ cells or supernatant p14 wre measured by flow cytometry or ELISA at 48h post-infection. (B) Schematic representation of HIV-infected T cells and pDC co-culture design. HIV-infected T cells for 48 hours were mixed with pDCs. IFN-I expression and HIV-1 replication in supernatants are measured 24h after co-culture. (C) CD4+ T cells infected with HIV-1 or HIV-1ΔVpr for 48 hours were co-cultured with or without purified autologous pDCs. HIV-1 replication was measured by p24 ELISA at 24 hour after co-culture. (D) Relative inhibition of replication of HIV-1 or HIV-1ΔVpr in CD4+ T cells by pDCs is summarized with three different donors. (E) IFN-α in the supernatant was measured at 24 hours after co-culture by ELISA. (F) Isotype control or α-IFNAR1 antibody was added at the beginning of co-culture, and HIV-1 replication (p24) was measured at 24 hour after co-culture. Error bars represent ±SD of triplicate experiments. *, ** and *** indicate p<0.05, <0.01 and <0.001, respectively.

### HIV-infected T cells activate pDC via CD4-dependent but RT-independent mechanisms

To define how HIV-infected T cells activate pDC in the coculture model, we found that sCD4 blocked IFN-I induction in a dose-dependent manner (Fig. 2A). Consistent with the IFNAR blockade results (Fig. 1F), sCD4 also elevated HIV-1 replication in the co-culture model and abrogated the Vpr activity to further enhance HIV-1 replication (Fig. 2B). In contrast, inhibition of RT (reverse transcription) in the coculture model showed no effect on IFN-I induction by T cells with established infection of either Vpr+ or Vpr-viruses (Fig. 2C). Consistent with predominantly established infection in T cells, RT inhibition showed only minor effect on HIV-1 replication, probably due to low levels of new infection (Fig. 2D).

**Figure 2.**
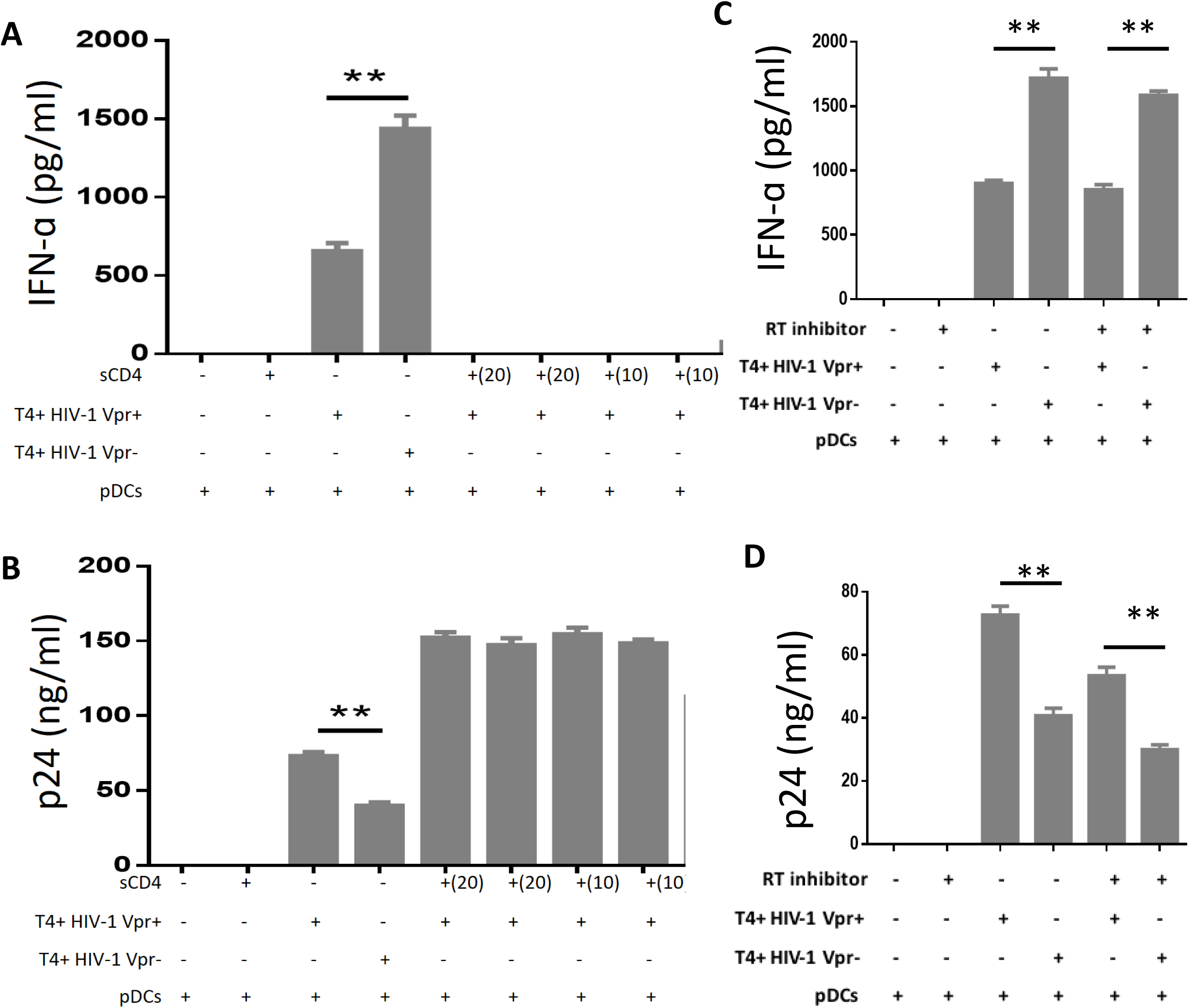
HIV-infected T cells activate pDCs via CD4-dependent but RT-independent mechanisms. (A) HIV-1 or HIV-1ΔVpr infected CD4+ T cells were cocultured with or without pDCs, with sCD4 at 20 μg/ml, 10 μg/ml or 2 μg/ml was added at the beginning of coculture. IFN- α in supernatants was measured at 24 hours after co-culture by ELISA. (B) p24 in supernatants was measured at 24 hours after co-culture by ELISA. (C) Activated primary CD4+ T-cells were infected with HIV-1 (MOI = 1) for 48 h. at the same time with viral infection. RT inhibitors nevirapine is 5 µm. p24 levels in the supernatants were quantified by ELISA. (D) HIV-1 or HIV-1ΔVpr infected CD4+ T cells were cocultured with or without pDCs. The reverse transcriptase (RT) inhibitor nevirapine (NVP) was added (5 µm) at the beginning of coculture. Supernatant IFN- α was measured at 24 hours after co-culture by ELISA. (D) Supernatant p24 was measured at 24 hours after co-culture by ELISA. Error bars represent ±SD of triplicate experiments.

### Virion-associated Vpr modulates IFN-I induction in pDCs via its TET2 degradation activity

HIV-1 activates pDCs through endosomal recognition of viral RNA by TLR7(Beignon et al., 2005; Lepelley et al., 2011). To test how Vpr in HIV-1 particles affects activation of pDCs, we utilized the TLR7/8 agonist, R848, to activate pDCs in the presence of HIV-1 or HIV-1ΔVpr viruses. As shown in Figure 3A, cell-free HIV-1 alone at the used dose failed to activate pDCs as previously reported (Bloch et al., 2014; Su et al., 2014). However, wild type HIV-1 but not HIV-1ΔVpr viruses significantly reduced IFN-α induction by R848. To confirm that virion-associated Vpr was sufficient to suppress IFN-I production in pDC during R848 treatment, we utilized virus-like particles containing Vpr (VLP_VPR_) or no Vpr (VLP) to treat pDCs (Fig. S2A). Similar to cell-free HIV-1 viruses, VLP +/- Vpr failed to activate pDC and pDC treated with VLP_VPR_ showed significantly reduced IFN-I expression in response to R848 (Figure 3B). When the kinetics of IFN-I mRNA induction was measured (Figure S2B), VLP_VPR_ impaired IFN-I expression during the early induction phase in response to R848. Virion-associated Vpr thus contributes to the reduced IFN-I induction in pDCs.

**Figure 3.**
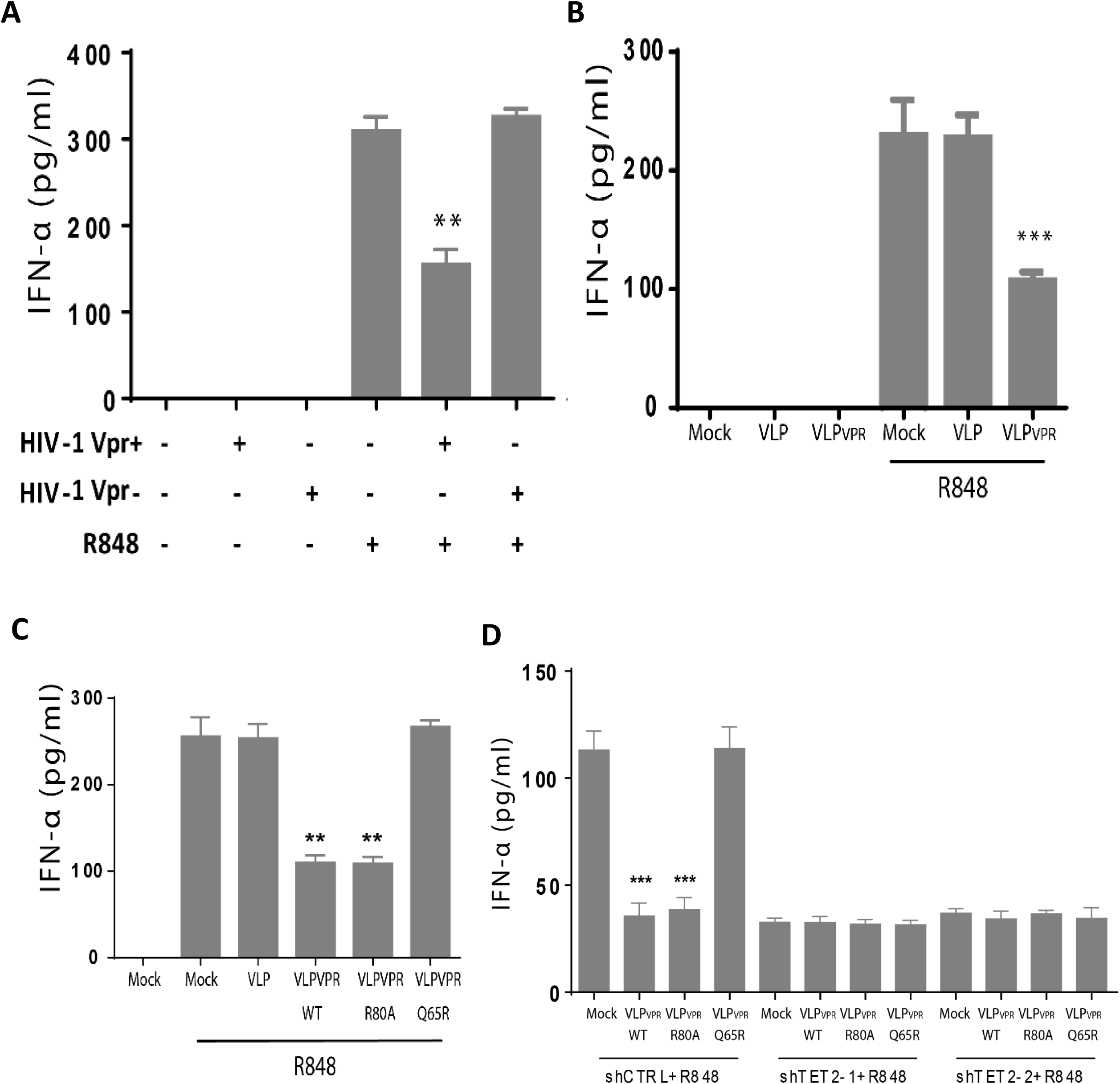
Virion-associated Vpr degrades TET2 to modulate IFN-I induction in pDCs. (A) pDCs were exposed to cell-free HIV-1 or HIV-1ΔVpr viruses for 10 hours, followed with R848 treatment for 16 hours. Supernatant IFN-α was measured by ELISA post R848 treatment. (B) pDCs were exposed to VLP or VLP_VPR_ in the absence or presence of R848. Supernatant IFN-α was measured at 16 hours by ELISA post R848 treatment. VLP or VLP_VPR_ pre-treat pDCs for 10 hours. Error bars represent ±SD of triplicate experiments. (C) pDCs were exposed to VLP_VPR-WT,_ VLP_VPR-R80A_ or VLP_VPR-Q65R_ for 10 hours and treated with R848. Supernatant IFN-α was measured at 16 hours post stimulation. (D) Human pDC-like cell line Gen2.2 cells were transduced with GFP-expressing shRNA control (shCTRL) or shRNA targeting TET2 (shTET2-1 and shTET2-2). TET2 knock-down efficiency was measured by real-time quantitative PCR and western blotting of the protein levels. Two independent TET2 shRNAs both stably reduced TET2 mRNA levels and protein levels in pDC cell line GEN2-2 (Fig. S3). GEN2-2 cells with shCTRL, shTET2-1 and shTET2-2 were exposed to R848 and VLP_VPR-WT,_ VLP_VPR-R80A_ or VLP_VPR-Q65R_. Supernatant IFN-α was measured at 24 hours post treatment. Error bars represent ±SD of triplicate experiments. ** and *** indicate p<0.01 and p<0.001, respectively.

Among a range of functions of the Vpr protein, Vpr-binding to VprBP and Vpr-dependent G2 arrest are the most extensively explored features (Belzile et al., 2007). We used Vpr mutants with specific defective activities of Vpr to define the Vpr-mediated impairment of IFN-I production in pDCs. Both Vpr_Q65R_ and Vpr_R80A_ mutants are defective for induction of G2/M arrest (Belzile et al., 2007). Vpr_Q65R,_ but not Vpr_R80A_ mutant, is defective for binding with VprBP (Le Rouzic et al., 2007). We recently discovered that HIV-1 Vpr targets the Tet methylcytosine dioxygenase 2 (TET2), a methylcytosine dioxygenase, via VprBP for degradation(Lv et al., 2018; Wang and Su, 2019). We produced virus-like particles containing Vpr_WT_, Vpr_Q65R_ or Vpr_R80A_ and all three Vpr proteins were efficiently incorporated into VLP (Figure S2B and(Lv et al., 2018; Wang and Su, 2019)). Similar to the Vpr activity in suppressing IFN-I expression in pDCs, VLP_VPR-WT_ and VLP_VPR-R80A_ but not VLP and VLP_VPR-Q65R_ degraded TET2 proteins in pDCs (Figure 3C and Fig. S3). VLP_VPR-WT_ and VLP_VPR-R80A_ showed similar TET2 degradation (Fig. S3A) and inhibition of IFN-I induction in pDCs as measured by IFN-α protein (Figure 3C) or mRNA levels of IFN-α and IFN-β (Figure S3B), whereas VLP_VPR-Q65R_ (like VLP with no Vpr) showed no activity. When TET2 was depleted by shRNA (Fig. S3C), induction of IFN-I by R848 was reduced by > 2-fold and Vpr failed to further reduce its induction (Fig. 3D). Therefore, Vpr interaction with VprBP to degrade TET2, but not its G2 cell cycle arrest activity, is critical for the Vpr-mediated inhibition of pDC activation.

### Vpr reduces IRF7 expression in pDCs

To determine if Vpr altered expression of the critical signaling molecules involved in IFN-I expression in pDCs, we measured their protein levels and discovered that IRF7 protein levels, but not other molecules in the pathway, were reduced by ∼40 % in the presence of VLP_VPR-WT_ and VLP_VPR-R80A_ (Fig. 4A). We further showed that VLP_VPR-WT_ and VLP_VPR-R80A_, but not VLP or VLP_VPR-Q65R_, reduced IRF7 mRNA levels in pDCs (Fig. 4B). When IRF7 mRNA was measured in control or TET2-KD pDC, TET2-KD reduced IRF7 expression and abrogated the Vpr activity to further reduce IRF7 expression (Fig. 4C). These results suggest that Vpr degrades TET2 to reduce IRF7 gene expression in pDCs to modulate IFN-I induction.

**Figure 4.**
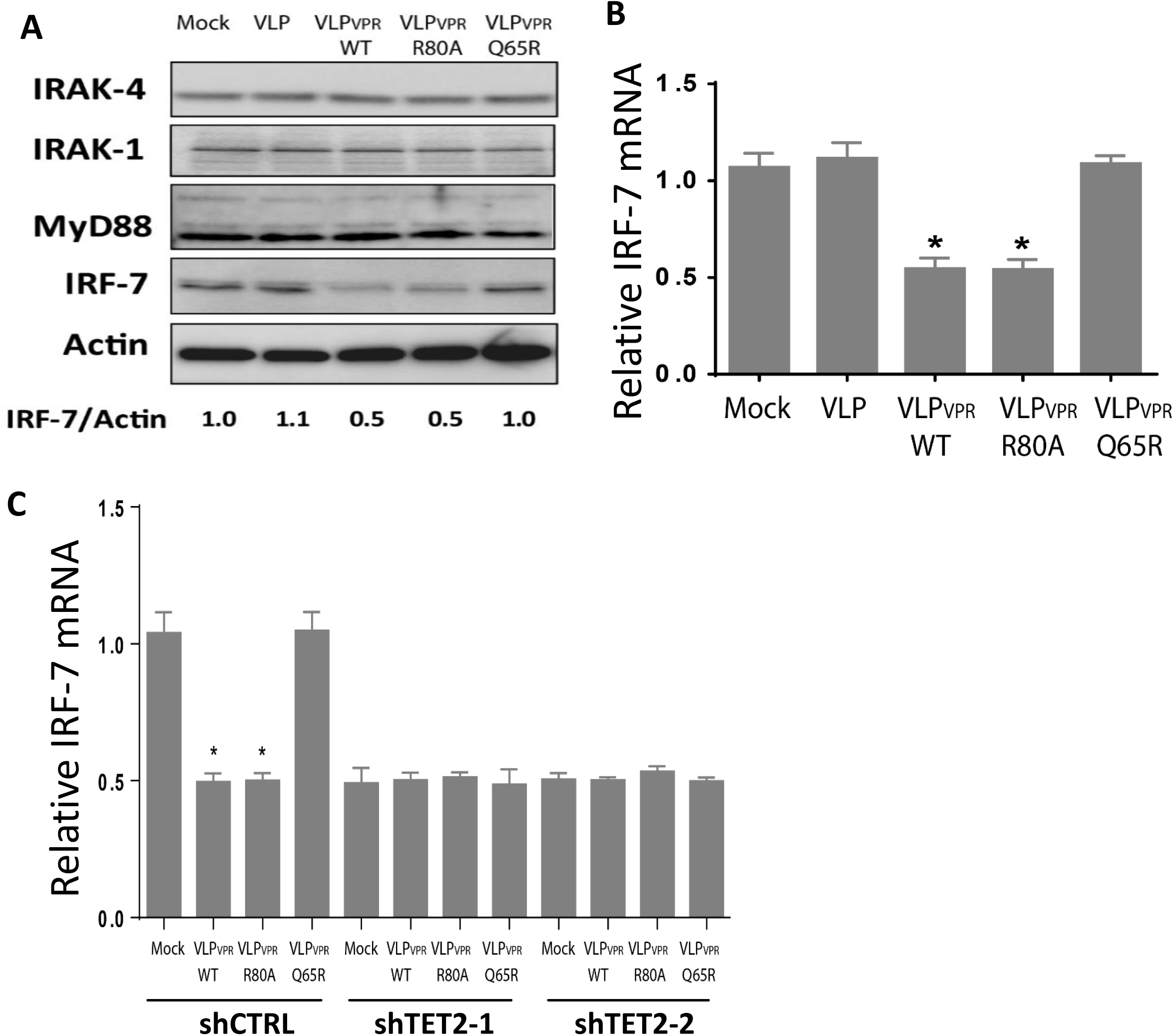
Vpr reduces IRF7 expression via TET2-dependent mechanisms in pDCs. (A) Primary pDCs were treated with VLP_VPR-WT,_ VLP_VPR-R80A_ and VLP_VPR-Q65R_ for 10 hours. Endogenous MyD88, IRAK-1/4, IRF7 and actin proteins were detected by Western blots. Primary pDCs were from three different donors were mixed for WB detection. (B) Primary pDCs were treated with mock, VLP, VLP_VPR-WT,_ VLP_VPR-R80A_ and VLP_VPR-Q65R_. IRF7 mRNA was measured at 10 hours after treatment. Error bars represent ±SD of triplicate experiments from three different donors. (C) GEN2-2 cells with shCTRL, shTET2-1 and shTET2-2 were treated with VLP_VPR-WT,_ VLP_VPR-R80A_ and VLP_VPR-Q65R_. IRF7 mRNA was measured at 24 hours after treatment. Error bars represent ±SD of triplicate experiments.

### Vpr degrades TET2 to prevent demethylation of the IRF7 promoter in pDCs

TET2 catalyzes the conversion of 5-methylcytosine (5mC) to 5-hydroxymethylcytosine (5hmC), leading to DNA demethylation to enhance gene expression (Abdel-Wahab et al., 2009). We showed that the pDC-specific CXXC5 protein recruited TET2 to the IRF7 promoter (Fig. 5A-C). To determine the 5hmC level on the IRF7 promoter in pDCs, we performed the anti-5hmC ChIP-qPCR assay on the IRF7 promoter and showed TET2 depended on CXXC5 to demethylate 5mC to 5hmC at the CpG island (Fig. 5D). Interestingly, correlated with their ability to degrade TET2, VLP_VPR-WT_ and VLP_VPR-R80A_ but not VLP_VPR-Q65R_ substantially reduced 5hmC levels at the IRF7 promoter in pDCs (Fig. 5E). These results demonstrate that HIV-1 Vpr reduced IRF7 and IFN-I gene expression through reducing TET2-dependent demethylation of the IRF7 promoter (Fig. 5F).

**Figure 5.**
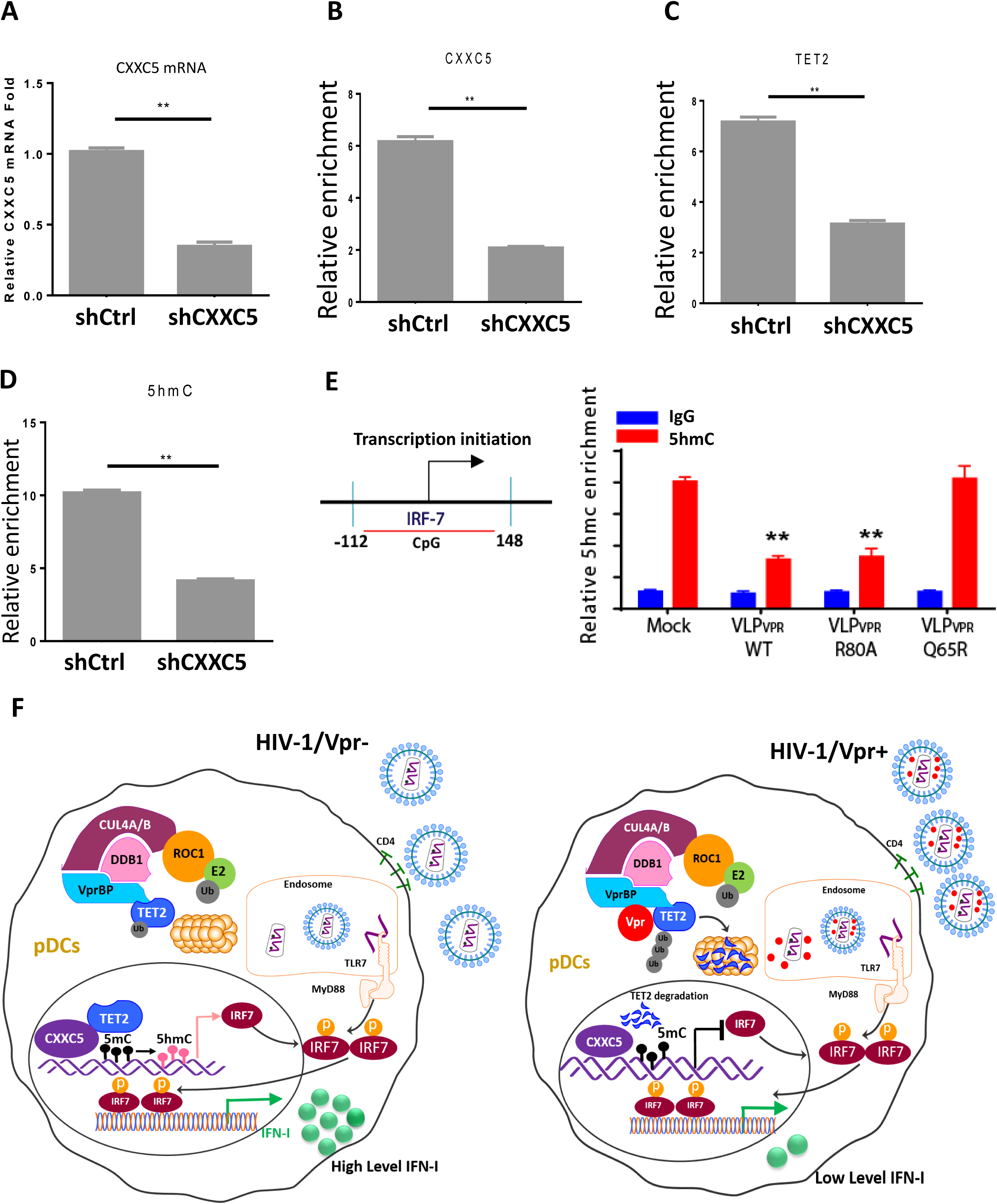
Vpr modulates IRF7 expression in pDCs via CXXC5/TET2-dependent demethylation of the IRF7 promoter. (A) Human pDC-like cell line Gen2.2 cells were transduced with control shRNA (shCTRL) or shRNA targeting CXXC5 (shCXXC5). CXXC5 knock-down efficiency was measured by real-time quantitative PCR. (B) ChIP assays were performed using antibodies against CXXC5 or isotype control IgG, followed by qPCR analysis of IRF7 promoter in shCTRL and shCXXC5 Gen2.2 cells. (C) ChIP assays were performed using antibodies against TET2 or isotype control IgG, followed by qPCR analysis of IRF7 promoter in shCTRL and shCXXC5-transduced Gen2.2 cells. (D) ChIP assays were performed using antibodies against 5hmC or isotype control IgG, followed by qPCR analysis of IRF7 promoter in shCTRL- and shCXXC5-Gen2.2 cells. (E) ChIP assays were performed using antibodies against 5hmC or isotype control IgG, followed by qPCR analysis of IRF7 promoter in the presence of VLP_VPR-WT,_ VLP_VPR-R80A_ or VLP_VPR-Q65R_. ChIP DNA level in each experiment was normalized to the IgG isotype control level (defined as 1). Error bars are derived from three technical replicates. (F) Schematic diagram of Vpr-mediated suppression of IRF7/IFN-I in pDCs to facilitate HIV-1 replication in CD4+ T cells. Upon endocytosis of HIV-1 virions in pDCs, the Vpr protein binds VprBP to target TET2 for degradation, leading to impaired demethylation of the IRF7 promoter and reduced IRF7 expression in pDCs. Reduction of IRF7 and IFN-I by Vpr in pDCs will facilitate HIV-1 replication in CD4+ T cells and other HIV-1 target cells.

## Discussion

Vpr is a highly conserved HIV-1 protein which is associated with high viral load and required for full pathogenesis by unclear mechanisms (Gibbs et al., 1995; Goh et al., 1998). We report here that Vpr promoted HIV-1 replication in CD4+ T cells when co-cultured with pDCs via IFN-I signaling dependent mechanisms. HIV-1 Vpr impaired IFN-I production in pDCs to confer partial resistance of HIV-infected-CD4+ T cells to resist viral inhibition by pDCs. Moreover, we showed that Vpr reduced IRF7 transcription and IFN-I production through degradation of TET2 in pDCs (Figure 5).

In agreement with a lack of IFN-I induction in T cells, Vpr didn’t affect HIV-1 replication in purified and activated CD4+ T cells (Figure S1B)(Balliet et al., 1994; Planelles et al., 1995). However, Vpr conferred HIV-infected CD4+ T cells to partially resist co-cultured autologous pDCs (Figure 1). As professional IFN-I-producing cells, pDCs rapidly produced high levels of IFN-I upon exposure to HIV-infected CD4+ T cells. We blocked IFN-I signaling and found HIV-1 replication were elevated to the same level in the presence or absence of Vpr when HIV-infected CD4+ T cells were co-cultured with pDCs (Fig. 1F), indicating that the major Vpr effect on pDC-mediated suppression of HIV-1 in T cells was due to differential induction of IFN-I. Interaction of HIV-infected T cells with CD4 was critical for the induction of IFN-I in pDC, although inhibition of reverse transcription showed no effect. Together the data indicate that pDC efficiently sensed HIV-infected T cells via CD4-depedent mechanism to induce IFN-I, which was impaired by Vpr.

IFN-I induces a number of anti-viral factors to inhibit HIV-1 replication in target cells (Kane et al., 2016), including APOBEC3, SAMHD1 and Mx2 (Hatziioannou and Bieniasz, 2011; Hotter and Kirchhoff, 2018). The results presented here showing that Vpr impaired IFN-I production in pDCs may partly account for Vpr-mediated enhancement of viral replication in SIV-infected monkeys and in HIV-1 infected humanized mice (Gibbs et al., 1995; Hoch et al., 1995)’ (Sato et al., 2013). Therefore, HIV-1 seems to have evolved to efficiently activate pDC via CD4-dependent endocytosis and TLR7 signaling. However, the magnitude of IFN-I induction is modulated by the Vpr protein to a low level that other HIV-1 accessory proteins are able to counteract the specific anti-HIV ISG restriction factors to maintain HIV-1 infection.

It is reported that Vpr may have a positive effect on IFN-I induction during HIV-1 infection of monocytes-derived DC (MDDCs) or monocytes-derived macrophages (MDMs) (Zahoor et al., 2015; Zahoor et al., 2014). In contrast, others have reported a suppressive role of Vpr on IFN-I production in MDDCs and MDMs (Harman et al., 2011; Mashiba et al., 2014), or in cell lines, such as HEK293, HeLa and HeLa-derived STING-expressing reporter cells (Vermeire et al., 2016).

Compared to conventional dendritic cells and macrophages, human pDCs express only the endoplasmic TLR7/9 but not other pattern recognition receptors (PRR) (Colonna et al., 2004). Once pDCs sense pathogens through TLR7/9, signaling is mediated via MyD88 (myeloid differentiation factor 88), a docking protein for IRAK1/4 (IL-1R-associated kinase 1/4). IFN-regulatory factor 7 (IRF7) is then phosphorylated and translocated into the nucleus to promote IFN-I gene expression (Honda et al., 2005b). It is well documented that pDCs uniquely express constitutive levels of IRF7 involving reduced methylation levels at the IRF7 promoter (Kulaeva et al., 2003). We further discovered that Vpr degraded TET2 through the CRL4^VprBP^ E3 ligase ligase (Lv et al., 2018) to suppress the IRF7 gene expression (Figure 4). CpG island on the human IRF7 promoter has been located around the TATA box spanning from −271 to +382 bp (Lu et al., 2000). We showed that Vpr-treated pDCs (or with TET2 shRNA) had reduced 5hmc levels at the IRF7 promoter (Figure 5E), associated with reduced IRF7 levels. Since TET2 does not have a specific DNA binding domain, we also showed that the pDC-specific factor CXXC5 was involved to recruit TET2 to demethylate the IRF7 promoter (Fig. 5 and (Ma et al., 2017)). It is likely that TET2 also demethylate other genes recruited by CXXC5 or other factors. Other putative target genes affected by Vpr or depletion of TET2 in pDCs are of great interest and will be studied in the future.

In summary, we have discovered that Vpr targets the TET2-IRF7 axis to modulate IFN-I induction in pDCs, leading to elevated HIV-1 replication in HIV-infected T cells. This Vpr activity depended on its interaction with the VprBP-CRL4 complex to degrade TET2. Interestingly, The constitutive expression of IRF7 and CXXC5 in pDCs is maintained by TET2-mediated demethylation of the IRF7 promoter. The dissection of the molecular mechanisms modulating IFN-I induction in pDCs by Vpr during HIV-1 infection could uncover new targets or strategy to treat HIV-1 replication and pathogenesis.

## Materials and Methods

### Viruses and plasmids

pNL4-3-R+ and pNL4-3-R- were obtained from the NIH AIDS Reagent program (Adachi et al., 1986; Connor et al., 1995; He et al., 1995). Flag-Vpr_Q65R_ and Flag-Vpr_R80A_ were generated by site-directed mutagenesis and verified by DNA sequencing.

### VLP stock production and transduction

For the production of virus-like-particles (VLP) carrying Vpr or Vpr-mutants, we transfected 293T with 6 μg p3 x FLAG-CMV-Vpr, 15 μg pMDLg/pRRE, 4 μg pRSV-Rev, and 5 μg pCMV-VSV-G. The VLP stock was treated with DNase (25 U DNase per 0.5 ml virus) before transduction. Concentration of VLP stocks were quantified by p24 ELISA assay (Frederick National Laboratory for Cancer Research – AIDS and Cancer Virus Program). All VLPs of 500 ng p24 amount and virus infections were performed by spin inoculation for 2 hours at 1500 g and 37 °C with 8 μg/ml polybrene (O’Doherty et al., 2000).

### Cell cultures and stimulation

pDC-like Gen2.2 cells (Chaperot et al., 2001) were kept as stock in our lab. Gen2.2 cells were maintained in RPMI 1640 medium containing 10% FBS, 1% Pen/Strep antibiotics, 2mM glutamine, 10mM HEPES, 1x nonessential amino acids and 1x Pyruvate (all from GIBCO). 293T cells were cultured in DMEM containing 10% FBS and 1X Antibiotic-Antimycotic (Invitrogen). Human buffy coats were obtained from the Gulf Coast Regional Blood Center. Primary human peripheral blood mononuclear cells (PBMCs) were isolated from buffy coats using Ficoll-paque gradient. BDCA-4 Microbead Kit (Miltenyi Biotec) was used to isolate human pDCs from PBMCs. Primary CD4+ T cells were negatively isolated from PBMCs using untouched dynabeads human CD4 T cell kit (Invitrogen, CAT# 11352D) according to manufacturer’s manual. Isolated CD4 T cells were stimulated with anti-CD3/anti-CD28 coated magnetic beads (Gibco, CAT#11131D) at bead:cell ratio 1:1, and maintained in RPMI 1640 (Invitrogen) supplemented with 10% fetal bovine serum (FBS) plus 200 IU/ml human recombinant IL-2 (NIH AIDS Reagent Program). Beads were removed at day 3 after the stimulation. 1 μg/ml R848 (Invivogen, tlrl-r848), 1μm CpG-A (Invivogen, tlrl-2216) and 1μm CpG-B (Invivogen, tlrl-2006) are used to activated pDCs. 10 μg/ml R848 was used to activate Gen2.2 cells.

### shRNA and RNA quantitation by real-time PCR

pGIPZ-lentiviral vectors expressing control (non-targeting), TET2-targeting shRNA were purchased from Dharmacon, GE Healthcare (Lafayette, CO). shRNA targeting human TET2 has mature antisense sequence: TAAGTAATACAATGTTCTT (clone ID: V3LHS_363201). Lentiviruses for shRNA targeting TET2 were produced by CaCl2-BES transfection of 293T cells (10 cm plate) using 10 ug vector, 15 μg pMDLg/pRRE, 4 μg pRSV-Rev, and 5 μg pCMV-VSV-G. Gen2.2 cells were transduced with ∼ 200 ng p24 of lentiviruses, and after 3 days sorted by GFP+ by flow cytometry. The recovered GFP+ cells were validated by western blot. For detection IFN-I and IRF7 expression. The primers are as following. IFN-a, F: GCCTCGCCCTTTGCTTTACT, R: CTGTGGGTCTCAGGGAGATCA. IFN-b, F: GTGCCTGGACCATAGTCAGAGTGG, R: TGTCCAGTCCCAGAGGCACAGG. IRF7, F: ACAGCACAGGGC GTTTTATC, R: TCCCGGCTAAGTTCGTACAC. GAPDH, F: CATGAGAAGTATGACAACAGCCT, R: AGTCCTTCCACGATACCAAAGT. Real-time PCR was performed using the SYBR green fluorescence system. IRF7 CHIP primer, F: GGGTTCCGCTGGTCGCATCC, R: CCCGCCCGACCCTCATCTCT.

### ELISA

ELISA kits for human IFN-α (Mabtech; 3425-1H-20) were used according to the manufacturer’s instructions. ELISA kits for p24 (Leidos, Biomedical Research Inc.) were used according to the instructions of the manufacturer.

### Antibodies

PerCP-Cy5-5-conjugated CD123 (Biolegend), FITC-conjugated DR (Biolegend), FITC-conjugated CD3 (Biolegend), APC-Cy7-conjugated CD4 (Biolegend), anti-5hmc (ACTIVE MOTIF) and anti-TET2 was obtained from Millipore (MABE462).

### Statistical analysis

All statistical analyses were performed with the unpaired Student’s t-test, and were considered significant when the p-value was less than 0.05. *, ** and *** indicate respectively p-values of less than 0.05, 0.01 and 0.001.

## Authorship contributions

Q.W. and L.T. designed and performed experiments, and analyzed the results and wrote the paper; L.L, Y.Z, L.C., H.Y. performed experiments and analyzed data; L.Z. and Y. X. provided reagents and analyzed data; L.S. conceived the research project, planned, designed the experiments, analyzed results and wrote the paper.

## Acknowledgements

We thank Liqun Chi and Yaxu Wu for technical assistance. We also thank the University of North Carolina (UNC) Division of Laboratory Medicine for animal care; University of North Carolina at Chapel Hill Center for AIDS Research (P30 AI50410), Flow Cytometry Core Facility, and the UNC Animal Histopathology Core Facility. We thank all members in the Su laboratory for critical reading and/or discussion of the manuscript.

We thank the NIH AIDS Research and Reference Reagent Program for kindly providing reagents. We thank Amanda Pons Taylor from Lineberger Comprehensive Cancer Center at the University of North Carolina at Chapel Hill for reading the manuscript. This study was supported in part by NIH grants AI095097 (to L.S.), CA163834 (to Y.X.) and AI127346 to L.S. and Y.X. The funders had no role in data analysis, preparation, or decision to publish the manuscript.

